# Vision Transformers Enable Advanced Plant Phenotyping in Controlled Environments

**DOI:** 10.64898/2026.09.04.748299

**Authors:** Janou Milligan, Anand Seethepalli, Aristeidis Tsaris, Xiao Wang, Larry York, John Lagergren

## Abstract

Reliable plant segmentation in high-throughput phenotyping must transfer across species and imaging conditions without repeated model tuning or extensive reannotation. We compare three segmentation strategies using images from Oak Ridge National Laboratory’s Advanced Plant Phenotyping Laboratory: (i) fixed color-based thresholding, (ii) supervised U-Nets trained from scratch, and (iii) pretrained vision transformers fine-tuned for binary segmentation. Models were evaluated on a held-out test set and a generalization set that comprised unseen species. On the held-out test set, thresholding, the best U-Net, and the best vision transformer achieved mean Dice scores of 58.3, 96.6, and 97.3, respectively. On the generalization set, the corresponding Dice scores were 56.5, 86.2, and 95.7. Thresholding remained effective on some datasets but failed when plant appearance changed. Supervised U-Net training resolved within-distribution errors but failed to generalize to novel species and backgrounds. Pretrained vision transformers consistently produced high-accuracy segmentations across the evaluated species, views, soil backgrounds, and tray types. These results benchmark the practical progression from fixed rules to task-specific supervision and pretrained visual representations for controlled-environment plant phenotyping.

## 1 Introduction

A central challenge in plant biology is connecting genes with phenotypes, which are determined by complex interactions between a plant and its environment [1, 2]. This interaction results in structural and phenotypic changes that vary widely between plant species and conditions. Addressing this challenge is important for developing crops and biological systems relevant to biotechnology, bioenergy, and emerging biological pathways to critical minerals [3, 4]. Plant phenotyping is a method that helps address this challenge, where large numbers of biologically meaningful plant traits measured across genetically diverse populations can be associated with genetic variation. However, this process traditionally relies on manual measurement of plant phenotypes (height, canopy architecture, greenness, etc.), which is labor intensive, expensive, low throughput, and often destructive to the plant [5]. Recent advancements in laboratory automation, robotic manipulation, and remote sensing have transformed this paradigm toward high-throughput, automated phenotyping in controlled greenhouse settings [3, 2, 6]. A persistent challenge in facility-scale phenotyping is that segmentation models trained for one experiment often fail when applied to new species, developmental stages, background artifacts, and treatment responses [6, 7]. This domain shift limits reuse of computer vision models and forces repeated annotation and retraining across experiments.

The Advanced Plant Phenotyping Laboratory (APPL; https://www.ornl.gov/appl) at Oak Ridge National Laboratory (ORNL) addresses many of the classical bottlenecks in plant phenotyping through automated, time-resolved, multimodal remote sensing in controlled environments [3]. APPL’s above-ground imaging modalities include two red-green-blue (RGB) sensors (side-view and top-view), two hyperspectral sensors (visible-to-near-infrared and shortwave-infrared), as well as a multispectral, infrared, chlorophyll fluorescence, and 3D-laser-scanning sensor [3, 8].

Although APPL collects multiple remote sensing modalities, RGB imagery remains a central modality because it is inexpensive, high resolution, broadly available across phenotyping systems, and commonly used to derive plant area and color traits. By repeatedly imaging plants over their development, APPL enables non-destructive measurement of dynamic traits such as projected biomass, canopy architecture, color, growth rate, and stress response across large experimental populations [3]. However, each experiment can generate tens to hundreds of thousands of images whose biological value depends on accurate segmentation of heterogeneous plant pixels from backgrounds across camera positions, species morphologies, and plant developmental stages. Segmentation errors propagate directly into down-stream trait estimates, biasing measurements of projected area, canopy structure, color indices, growth dynamics, and stress responses used for genotype-phenotype association [5]. As a result, segmentation is a critical initial step in APPL image-based phenotyping workflows.

Fixed color-based thresholding is a viable approach for controlled-environment plant phenotyping because it is fast, interpretable, and requires no labeled training data [6, 7]. However, the performance of such approaches depends on high-contrast separation between plant tissue and the surrounding background. Fixed pixel-level thresholding rules may therefore degrade as plant pigmentation, developmental stage, senescence, illumination, soil, pots, or tray materials change. Supervised convolutional neural networks, such as U-Net [9], replace fixed rules with task-specific spatial and contextual features that are learned from manually annotated masks and have become a widely used approach for plant segmentation. However, models trained on one experiment may not generalize to changes in species, plant architecture, or background [7]. Fine-tuning pretrained vision transformers (ViTs) [10, 11, 12] provides a third strategy by adapting visual representations learned from large external datasets to plant segmentation using the available domain-specific annotations.

This work evaluates three plant-segmentation paradigms spanning alternate assumptions and supervision requirements: (i) a fixed color-threshold method with no learned parameters, (ii) U-Net models trained from scratch using task-specific annotations, and (iii) pretrained ViT encoders fine-tuned using the same annotations. The learned models were trained using 1,500 manually annotated RGB images of poplar (*Populus trichocarpa*) and switchgrass (*Panicum virgatum*) and a validation set of 250 images. All three strategies were evaluated on a test set of 250 held-out images of poplar and switchgrass and on 1,000 images from arabidopsis, eucalyptus, pennycress, sorghum, and soybean, spanning APPL above-ground RGB modalities with varying soil backgrounds and tray types. This approach was designed to evaluate the utility of fixed rules, task-specific supervised learning, and pretrained visual representations under facility-relevant domain shifts, rather than isolating the effects of convolutional and transformer-based neural network architectures with and without pretraining. Model scale, window size, patch size, and inference strategies for learned models were additionally evaluated. By testing these approaches and their generalization across species, tray types, and backgrounds, this study assesses whether pretrained vision transformers provide a practical route toward reusable segmentation models for controlled-environment phenotyping.

## 2 Methods

### 2.1 Data

All image data used in this work were collected at ORNL’s APPL robotic greenhouse system, designed by Photon Systems Instruments (PSI) [13, 14], which supports up to 520 trays containing one or more plants for experiments lasting days to months [15, 3]. APPL is a temperature-controlled facility that automatically applies watering, nutrient, temperature, and lighting schedules to simulate real-world environments. Each plant travels on a robotic conveyor system through an array of imaging stations spanning top-view and side-view RGB, hyperspectral, multispectral, infrared, chlorophyll fluorescence, and 3D-laser-scanning modalities for above- and below-ground phenotyping. APPL combines these remote-sensing systems to collect time-resolved, multimodal images of plants growing under simulated environmental conditions. The two biomass feedstocks considered primarily in this work are poplar and switchgrass, each with population-scale genome annotations and decades of research connecting their genomes to phenotypes [16, 17, 5, 18]. In addition, arabidopsis, eucalyptus, pennycress, sorghum, and soybean were also used to evaluate model generalization and robustness to novel species outside the training datasets. See Table 1 for a summary of the dataset composition with further details given below.

**Table 1:** Dataset composition. Training, validation, and test samples were drawn from poplar and switchgrass datasets, while additional species were reserved exclusively for generalization evaluation. RGB1 corresponds to APPL’s side-view RGB sensor and RGB2 to APPL’s top-view RGB sensor. The total counts are 1,500 training, 250 validation, 250 testing, and 1,000 generalization images. All images and masks are publicly available upon publication.

| Species | Modality | Train | Validation | Test | Generalization |
| --- | --- | --- | --- | --- | --- |
| Poplar | RGB1 | 369 | 70 | 61 | - |
| Poplar | RGB2 | 369 | 70 | 61 | - |
| Switchgrass | RGB1 | 381 | 55 | 64 | - |
| Switchgrass | RGB2 | 381 | 55 | 64 | - |
| Sorghum | RGB1 | - | - | - | 100 |
| Sorghum | RGB2 | - | - | - | 100 |
| Sorghum-Rhizobox | RGB1 | - | - | - | 100 |
| Sorghum-Rhizobox | RGB2 | - | - | - | 100 |
| Soybean | RGB1 | - | - | - | 100 |
| Soybean | RGB2 | - | - | - | 100 |
| Eucalyptus | RGB1 | - | - | - | 100 |
| Eucalyptus | RGB2 | - | - | - | 100 |
| Arabidopsis | RGB2 | - | - | - | 100 |
| Pennycress | RGB2 | - | - | - | 100 |

Model development focused on APPL RGB imagery because it is widely used in plant phenotyping and provides the highest spatial resolution among APPL’s imaging modalities, making it particularly important for estimating plant morphology. During data acquisition, each plant was automatically imaged using APPL’s side-view (RGB1) and top-view (RGB2) sensors, providing paired, complementary views of the same plant. From the total dataset of APPL RGB images, 2,000 images were selected for manual annotation: 1,000 for poplar and 1,000 for switchgrass. For each species, 500 side-view images and their 500 paired top-view images were included. To ensure the training dataset captured phenotypic variance across APPL’s existing experiments, coarse masks for the side-view modality were generated using the fixed threshold pipeline described in Section 2. The images and masks were then used to extract shape-based features (e.g., height, area, and perimeter), shape-based indices (e.g., solidity, roundness, and eccentricity), and color-based features (e.g., maximum, minimum, and mean red, green, and blue). Principal component analysis (PCA) was used to reduce the dimensionality of the feature space to three principal components that captured the majority of the variance across plants and time points. Each principal component was used to partition the dataset into five sections. Then, latin hypercube sampling (LHS) [19] was used to randomly select four samples from each combination of principal component partitions (5^3^ = 125 total), resulting in 500 side-view RGB images per species. The corresponding top-view image acquired from each plant was additionally included, resulting in 500 images per species/modality combination (2,000 in total). The coarse masks and their derived features were used solely for sampling and were not used as training or evaluation labels. All reported segmentation scores were computed against manually annotated ground truth masks, which are described further below.

The training dataset was partitioned into 1,500 training images (75%), 250 validation images (12.5%) and 250 testing images (12.5%) splits. To prevent data leakage, each plant was assigned to a given split using a group-wise splitting method. A “group” was defined as all modalities and time points belonging to a given plant (defined using APPL’s `PlantID` metadata field). From the 2,000 samples, groups were randomly selected and their corresponding images added to the test, then validation, and finally the train split. Group selections were randomly resampled until the train/validation/test proportions matched 1500/250/250 images, respectively. Each split was confirmed to be disjoint from each other as a quality control.

To generate a generalization dataset, 1,000 images were curated across APPL experiments for five species: arabidopsis, pennycress, eucalyptus, soybean and sorghum. Available modalities included both side-view and top-view RGB for eucalyptus, soybean, and sorghum, and only top-view RGB for arabidopsis and pennycress. Sorghum grown in rhizobox trays (used for below-ground imaging) were also included to test model performance with new tray types. Due to a smaller number of available images compared to poplar and switchgrass, these images were manually selected. For each combination of species and modality/tray type excluding eucalyptus, arabidopsis, and pennycress, ten plant time series with ten time points each were chosen. The ten images per plant were sampled evenly from start to end of each available time series. Eucalyptus only had two available time points per time series and therefore included 50 plants with two time points each. Arabidopsis and Pennycress had variability in the number of time points per time series per experiment, resulting in 27 time series for pennycress and 38 for arabidopsis. Each time series was manually compared with all others in a given combination to include as much variation as possible. This strategy resulted in a dataset of 1,000 images (100 per combination) spanning different species, camera angles, experimental conditions, tray types, and growth stages.

All selected images were manually annotated for binary plant/background segmentation through CVAT Labeling Services [20]. The resulting masks were visually reviewed by scientists with expertise in plant phenotyping and computer vision before inclusion in the dataset. Quality control focused on plant omissions, inclusion of non-plant structures, and boundary errors. Observed systematic errors were reported to CVAT, which then corrected the identified subsets of images. The reviewed masks served as ground truth annotations for model training, validation, and evaluation.

### 2.3 Segmentation

This work compared (i) APPL’s fixed color-threshold pipeline, (ii) supervised U-Net models trained from scratch, and (iii) segmentation models based on pretrained ViT encoders. The threshold pipeline required no learned parameters. U-Net and ViT models were implemented in Python (3.12) using PyTorch (`torch`; 2.8.0) [21], with Segmentation Models PyTorch (`smp`; 0.5.0) [22] providing the U-Net architectures and PyTorch Image Models (`timm`; 1.0.20) [23] providing the ViT architectures and pretrained checkpoints. Each learned model input a 3-channel image tensor (*X* ∈ ℝ^*B*×3×*H*×*W*^) and output a 2-channel tensor of class logits (*Y* ∈ ℝ^*B*×2×*H*×*W*^), where *B* represents the batch size and *H* and *W* represent the height and width, respectively, of the input/output window. Since each RGB image is large, side-view RGB1: (6556, 4104, 3) pixels and top-view RGB2: (3006, 4104, 3) pixels, they were divided into smaller image tiles for training and inference. The threshold-based approach directly segmented each image into a 1-channel binary matrix (*Y* ∈ ℝ^*H*×*W*^) and did not require tiling. See Table 2 for a list of learned model variants and their parameter counts. Each model is described in further detail below.

**Table 2:** Model sizes. Parameter counts are reported for each learned segmentation model and backbone, including total parameters across the encoder and decoder. (a) shows the U-Net models with four ResNet backbones and two input sizes, while (b) shows the ViT models with two patch sizes and two input sizes.

| (a) U-Net |  |  | (b) ViT |  |  |  |
| --- | --- | --- | --- | --- | --- | --- |
| Backbone | Input Size | Parameters | Backbone | Patch | Input Size | Parameters |
| ResNet-34 | 224 | 24.4M | ViT-Small | 16 | 224 | 24.3M |
| ResNet-34 | 448 | 24.4M | ViT-Small | 16 | 448 | 24.5M |
| ResNet-50 | 224 | 32.5M | ViT-Small | 8 | 224 | 24.3M |
| ResNet-50 | 448 | 32.5M | ViT-Small | 8 | 448 | 25.2M |
| ResNet-101 | 224 | 51.5M | ViT-Base | 16 | 224 | 96.4M |
| ResNet-101 | 448 | 51.5M | ViT-Base | 16 | 448 | 96.8M |
| ResNet-152 | 224 | 67.2M | ViT-Base | 8 | 224 | 96.3M |
| ResNet-152 | 448 | 67.2M | ViT-Base | 8 | 448 | 98.1M |

To represent a classical non-learning segmentation workflow, we evaluated APPL’s fixed color-threshold pipeline on every test and generalization image. RGB channels were normalized to [0,1], and pixels satisfying

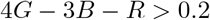

were classified as plant, where R, G, and B represent the red, green, and blue channels, respectively [24]. Images were pre-processed using a 5-pixel median filter before thresholding, and the resulting masks were post-processed by removing objects smaller than 150 pixels and filling holes smaller than 50 pixels. The threshold and all filter parameters were fixed before evaluation and applied unchanged across species, camera views, tray types, and dataset splits. The method further operated directly on full-resolution images, did not use learned parameters or tile-based reconstruction, and did not select or adjust parameters using the validation, test, or generalization data.

For the supervised baseline, a U-Net segmentation model [9] was implemented using the Segmentation Models PyTorch library (`segmentation_models_pytorch`). U-Net follows an encoder–decoder architecture in which the encoder progressively reduces spatial resolution while increasing feature depth, and the decoder restores spatial resolution using learned upsampling operations. Skip connections pass intermediate encoder features directly to corresponding decoder stages, preserving fine-scale spatial information that is often lost during downsampling. This structure is well suited for semantic segmentation because it combines high-level contextual features with local edge and shape information needed for pixel-level prediction, and is widely used in segmentation and plant phenotyping [5, 25]. In this work, the U-Net used Residual Network (ResNet) [26] encoder backbones selected from `resnet34`, `resnet50`, `resnet101`, and `resnet152`, with three-channel RGB inputs and two output classes. Unlike the ViT models, the U-Net baselines were trained fully supervised without using pretrained encoder weights. This design choice provided a comparison between segmentation strategies: conventional CNN-based segmentation trained from scratch and pretrained transformer-based segmentation under the same data splits, tile sizes, loss function, and optimization procedure (see Section 2.3 for training details). Accordingly, this comparison evaluates a practical progression from fixed thresholding, to supervised U-Nets trained from scratch, to fine-tuning pretrained visual representations, rather than whether the ViT architecture is inherently superior to U-Net. See Figure 1a for an illustration of the U-Net baseline model.

**Figure 1:**
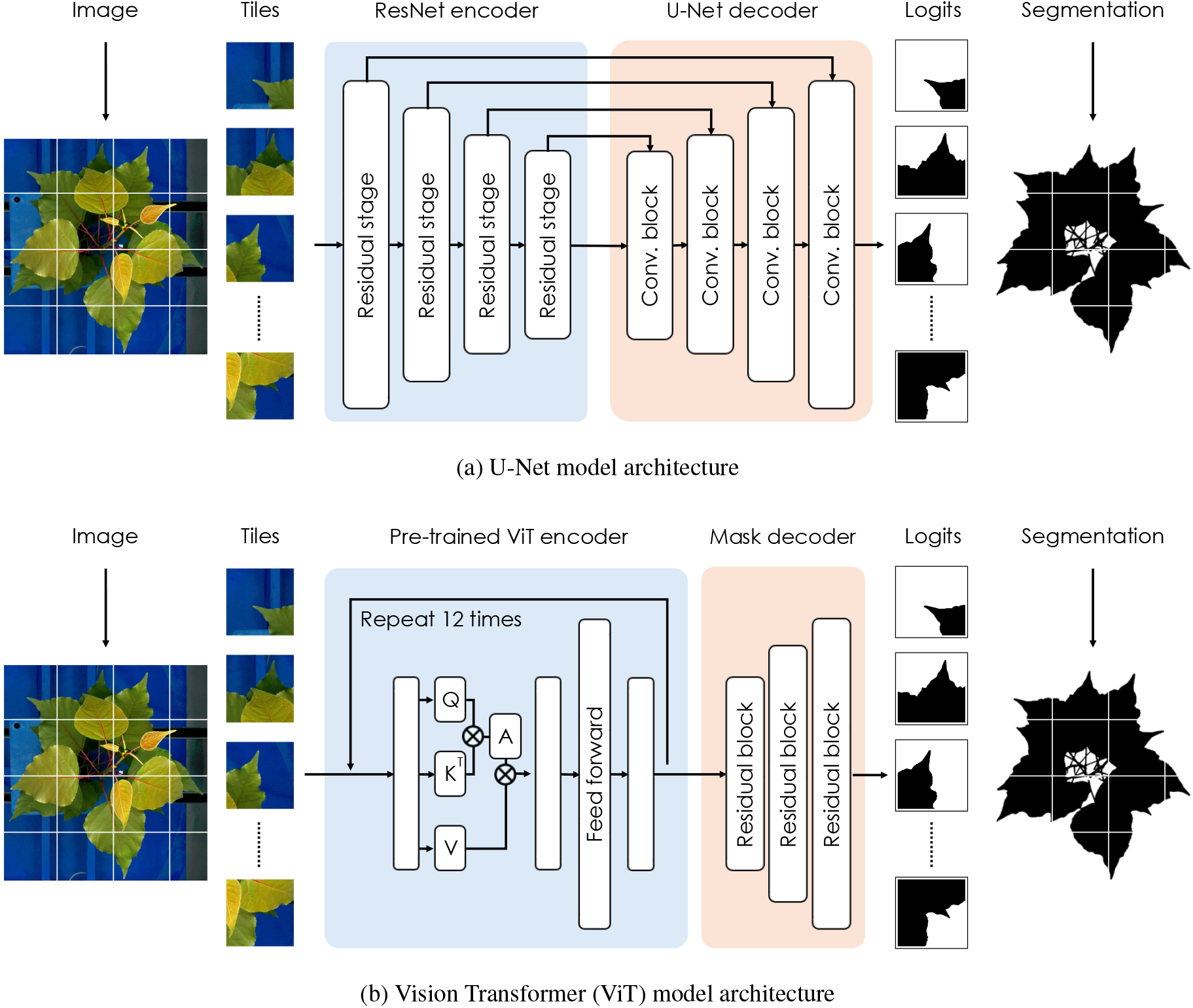
Segmentation model architectures. Images are divided into smaller tiles to accommodate the input size of each model. (a) Image tiles are processed by a ResNet encoder to extract hierarchical convolutional feature representations. A U-Net decoder combines intermediate decoder representations with internal convolution layers through skip connections and upsampling to produce dense pixel-level predictions for each tile. (b) Image tiles are processed by a pretrained transformer encoder to generate contextual feature representations. A lightweight convolutional decoder progressively upsamples the encoder output to produce dense pixel-level predictions for each tile. *Q, K, V*, and *A* represent the queries, keys, values, and attention weights in each multi-head self-attention layer, respectively. The feed forward block represents a simple multilayer perceptron and residual blocks represent 2 × upsampling and residual layers. Both models process RGB tiles and output dense pixel-level predictions for each tile.

For ViT-based segmentation, each RGB tile was encoded using a pretrained vision transformer backbone from the PyTorch Image Models library (`timm`). In the ViT architecture, image tiles were first partitioned into non-overlapping patches of size *P* ∈ *{*8, 16} pixels, producing an output spatial feature map of resolution *H/P* × *W/P* with channel dimension determined by the selected ViT backbone (ViT-Small or ViT-Base) [10]. Note the difference between image tiles and patches: original RGB1 and RGB2 images were divided into *H* × *W* image tiles, and each tile was further divided into 8 × 8 or 16 × 16 patches (image tokens) that were then passed to the ViT encoder. Both backbones contained 12 transformer layers, but ViT-Small used a 384-dimensional embedding with 6 attention heads per layer, whereas ViT-Base used a 768-dimensional embedding with 12 attention heads per layer, resulting in more parameters (Table 2). Only the final encoder feature map was used for decoding to the original tile resolution. To recover dense pixel-level predictions, a lightweight convolutional decoder progressively upsampled the encoded representation by a factor of two until the original input resolution was restored. Each upsampling stage consisted of nearest-neighbor upsampling, batch normalization, ReLU activation, a 3 × 3 convolution that reduced the channel dimension by half, and a two-layer residual convolutional block. The number of decoder stages was set to log_2_(*P*), yielding three stages for patch-8 models and four stages for patch-16 models. A final batch normalization, ReLU activation, and 1 × 1 convolution projected the decoder features to two output logits corresponding to background and plant classes. To account for different input sizes, the position embeddings were interpolated from the native 224 resolution to 448. See Figure 1b for an illustration of the ViT segmentation model.

Four pretrained ViT checkpoints were obtained from Hugging Face `timm`. ViT-Small with 8 × 8 patch size was initialized from `timm/vit_small_patch8_224.dino` [27], which used DINO self-supervised pre-training [11] on ImageNet-1k [28]. The other three models included ViT-Small with 16 × 16 patch size initialized from `timm/vit_small_patch16_224.augreg_in21k_ft_in1k` [29] and ViT-Base 8 × 8 and 16 × 16 patch size initialized from `timm/vit_base_patch8_224.augreg2_in21k_ft_in1k` [30] and `timm/vit_base_patch16_224.augreg2_in21k_ft_in1k` [31], respectively. These three checkpoints were pre-trained on ImageNet-21k and subsequently fine-tuned on ImageNet-1k. The ViT-Small with 8 × 8 patch size checkpoint was selected because no corresponding ImageNet-21k checkpoint was listed in the `timm` 1.0.20 model registry used for this work.

### 2.3 Training

All learned models were trained using the same data splits, tile sizes, loss function, optimization schedule, and validation procedure to align training and evaluation conditions for the U-Net and pretrained ViT strategies. Each model was trained on a single NVIDIA A100 GPU with 40 GB of memory. Training was performed on the 1,500-image poplar and switchgrass training split, with model checkpoints based only on the 250-image validation split. Both ViT and U-Net models were trained for binary semantic segmentation with two output classes, corresponding to background (0) and plant pixels (1). For each model backbone in Table 2, separate models were trained using tile sizes of 224 and 448 pixels in order to pair comparisons across model architecture family, scale, and input resolution.

Because the original RGB images were substantially larger than the model input windows and available GPU memory, training was performed on cropped image tiles rather than full-resolution images. For each training image, tile center coordinates were sampled from precomputed foreground and background “regions of interest” generated using a Euclidean distance transform of the inverse ground-truth mask. In particular, foreground coordinates satisfied *d < T* and background coordinates satisfied *d* ≥ *T*, where *d* denotes Euclidean distance and *T* denotes tile size in pixels. Foreground tiles were therefore drawn from plant regions, while background tiles contained only background pixels. Coordinates within *T/*2 pixels of image boundaries were excluded to ensure all sampled tiles were contained within the image. During training, 10 tiles were randomly sampled per image per epoch, producing 15,000 training tiles per epoch. To reduce class imbalance, tiles were randomly selected from either the foreground or background coordinate pool with equal probability when both pools were available. This produced an approximately 50/50 split between tiles containing plant pixels or only background.

Training tiles were randomly augmented during data loading using geometric and photometric transformations. Each tile was randomly horizontally flipped, randomly transformed with affine scaling and shear, and randomly adjusted for brightness and contrast. These augmentations were intended to expose each model to subtle changes in plant position, apparent size, viewing angle, and illumination to simulate variation in greenhouse conditions while preserving the semantic structure of the segmentation masks. After augmentation, all RGB tiles were normalized using ImageNet [28] channel mean, `[0.485, 0.456, 0.406]`, and standard deviation, `[0.229, 0.224, 0.225]`, before conversion to PyTorch tensors and transfer from CPU to GPU. Validation tiles were not augmented and were processed only with the same ImageNet normalization to ensure that validation performance reflected model generalization rather than stochastic transformations.

Models were optimized using combined cross-entropy and Dice loss. Cross-entropy loss provides pixel-wise supervision for the two-class background/plant classification problem, while Dice loss directly penalizes errors in overlap between predicted and ground-truth masks. The final objective was the equally weighted sum of the two terms:

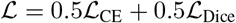

where the cross-entropy term, ℒ_CE_, takes the form:

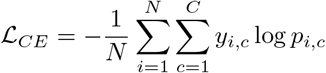

and the Dice term, ℒ_Dice_, takes the form:

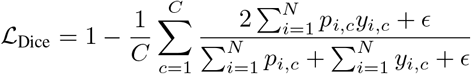

where *N* is the number of pixels being evaluated, *C* = 2 is the number of classes (background and plant), *i* indexes pixels, and *c* indexes classes. The term *y*_*i,c*_ is the one-hot encoded ground-truth label for pixel *i* and class *c*, while *p*_*i,c*_ is the predicted softmax probability assigned to class *c* at pixel *i. ϵ* = 10^−6^ was a constant to prevent division by zero. Cross-entropy loss was averaged over pixels in each batch, while Dice was computed per tile and class and then averaged over the batch. This combination of loss terms was used because APPL RGB images contain large background regions relative to plant area, making overlap-based optimization useful for reducing the effects of class-imbalance during training [32].

All models were trained with the AdamW [33] optimizer using a weight decay of 10^−4^. Training used automatic mixed precision and gradient accumulation to fit larger effective batch sizes in GPU memory. The micro-batch size was 14 tiles, and gradients were accumulated over nine micro-batches, giving an effective batch size of 126 tiles. U-Net models used a single learning rate for all trainable parameters. For ViT models, the pretrained encoder and randomly initialized decoder were optimized as separate parameter groups, with the encoder learning rate scaled by a factor of 0.1 relative to the decoder to adapt the pretrained backbone more conservatively. The learning rate, *η*, was scheduled using a linear warmup from zero to maximum learning rate, *η*_max_ for 10 epochs, followed by cosine decay of 50 epochs to the minimum learning rate, *η*_min_, and kept at the minimum learning rate for the remainder of training:

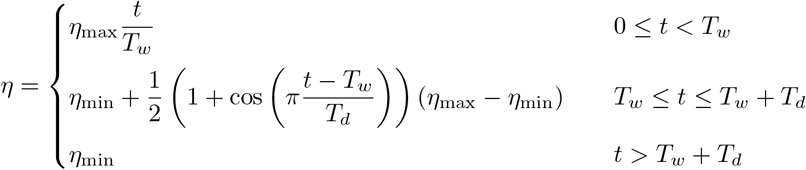

where, *t* denotes the training epoch, *T*_*w*_ = 10 is the warmup period, *T*_*d*_ = 50 is the cosine decay period, *η*_max_ = 10^−4^ is the maximum decoder learning rate, and *η*_min_ = 10^−6^ is the minimum learning rate. For ViT models, the same schedule was applied to both parameter groups, with the encoder maximum learning rate multiplied by 0.1 and both encoder and decoder sharing the same minimum learning rate. Each model (i.e., each ViT and U-Net variant) was trained for 100 epochs. See Supplementary Figure 1 for a visualization of the learning rate schedule.

After each epoch, models were evaluated on the full validation split using deterministic, non-overlapping tile grids. Validation tiles were generated across each image without random sampling or augmentation, and predictions from all tiles for a given image were concatenated before computing image-level Dice [34]. This image-level validation metric avoided over-weighting background-dominated images in APPL. Training and validation cross-entropy loss, Dice loss, total loss, and Dice score were logged after every epoch. The checkpoint with the highest mean validation Dice was saved for downstream testing and generalization evaluation.

### 2.4 Inference

The threshold baseline was applied directly to each full-resolution image using fixed image preprocessing and mask post-processing rules. The learned U-Net and ViT models used a tile-based algorithm described below. All three methods were scored using the same image-level Dice calculation against the manual ground truth masks. U-Net and ViT model inference was performed using the checkpoint with the highest mean validation Dice for each model backbone and tile-size configuration. Each trained model was applied to the full train, validation, test, and generalization splits to produce image-level segmentation masks and evaluation metrics. This procedure allowed downstream comparisons to use the same reconstruction and scoring pipeline across ViT and U-Net models. Because APPL RGB images are much larger than the model input size, each full-resolution image was segmented using a sliding-window tiling strategy. Images were normalized using the same ImageNet channel means and standard deviations used during training, then divided into tiles matching the model’s trained input resolution (224 × 224, or 448 × 448). Tile coordinates were generated with a stride of half the input resolution, producing a 50% overlap between adjacent tiles. Since the tile sizes did not divide the image resolution exactly, additional tiles were placed along the bottom and right image boundaries to ensure image coverage at the image boundaries.

For each tile, the models produced two-channel logits corresponding to background and plant classes, which were converted to class probabilities using softmax. Because each tile was segmented independently, predictions near tile boundaries had access to less surrounding image context than predictions near tile centers, often producing visible boundary artifacts. To reduce these artifacts, overlapping tile predictions were merged in probability space rather than mask space. Each tile probability map was weighted by a two-dimensional (2D) Hann window [35] before accumulation into a full-image probability map, and each pixel was normalized by the sum of weights contributing to that location. Weights were clipped to 10^−6^ to prevent division by zero. This weighting therefore emphasized predictions near tile centers while reducing the influence of less reliable predictions near tile boundaries. The final segmentation mask was obtained by assigning each pixel to the class with maximum normalized probability. The Hann-weighted overlapping inference described above was compared against classical non-overlapping inference, in which images were divided into non-overlapping tiles for prediction and corresponding masks concatenated together to full image resolution. Results are reported in Section 3 and the tile-based inference is illustrated in Supplementary Figure 2.

Segmentation accuracy was evaluated at the image level using Dice overlap between the reconstructed full-resolution prediction and the corresponding manually annotated mask. For each image, Dice was computed after full-image concatenation/aggregation so that the metric reflected the final mask used for phenotyping rather than tile-level performance. Inference time was also recorded for each image. No morphological post-processing, object filtering, or hole filling was applied to U-Net or ViT predictions. The threshold baseline used fixed median filtering, object removal, and hole filling as part of the standard APPL RGB image-processing pipeline. Dice overlap was reported as a percentage, with higher values indicating greater overlap between predicted and ground-truth masks. Mean and standard deviation of Dice were reported across each evaluation dataset. Standard deviation was chosen to characterize variation in image-level segmentation performance across plants and imaging conditions.

## 3 Results

Across both evaluation datasets, segmentation accuracy improved progressively from fixed color thresholding to supervised U-Net training and pretrained ViT fine-tuning (Table 3). The threshold-based method produced similar mean Dice scores on the test and generalization datasets but exhibited substantial variability, indicating that its performance depended strongly on plant appearance and imaging context. Supervised U-Nets largely resolved these errors for the test dataset but showed reduced accuracy and greater variability when applied to unseen species and backgrounds in the generalization set. Pretrained ViTs achieved the highest accuracy in both settings, with only a modest improvement over U-Net within distribution but a substantially larger and more consistent improvement on the multi-species generalization dataset. Validation loss and Dice convergence between learned models is shown in Supplementary Figure 3. The following sections examine the differences between models in greater detail, including the effects of model configuration, spatial resolution, failure modes, and tile-based inference.

**Table 3:** Segmentation accuracy. Image-level Dice scores (mean ± standard deviation, percentage points) are reported for the best-performing threshold-based, supervised U-Net, and pretrained ViT segmentation strategies on the within-distribution test set and the multi-species generalization dataset. Higher Dice values indicate greater overlap between predicted and manually annotated segmentation masks. Best-performing values for each evaluation dataset are shown in bold.

| Method | Training | Test Dice | Generalization Dice |
| --- | --- | --- | --- |
| Threshold | Fixed rule; no training | $58.3 \pm 28.5$ | $56.5 \pm 32.2$ |
| Best U-Net | Supervised from scratch | $96.6 \pm 4.0$ | $86.2 \pm 23.9$ |
| Best ViT | Pretrained and fine-tuned | **$97.3 \pm 3.5$** | **$95.7 \pm 10.1$** |

### 3.1 Within-distribution accuracy

Within-distribution evaluation revealed that the largest increase in segmentation accuracy came from replacing fixed color rules with learned segmentation. The threshold baseline achieved an overall image-level Dice score of 58.3 ± 28.5, with clear dependence on species and camera view (e.g., performance ranged from 86.5 ± 9.0 on poplar to 26.1 ± 23.0 on switchgrass top-view RGB2 images). The best performing U-Net, based on a ResNet-34 encoder with 448-pixel tiles, increased overall Dice to 96.6 ± 4.0, with a gain of 38.3 percentage points over thresholding, showing that task-specific supervision largely overcame failures of the fixed color rule. Pretrained ViTs provided a smaller but consistent improvement on species represented during training, reaching 97.3 ± 3.5 Dice, approximately 0.7 percentage points above the best U-Net with reduced standard deviation. Learned model performance was uniformly high for both poplar imaging modalities, where several ViT models exceeded 99% Dice on RGB2 images, while the largest performance differences between architectures occurred for switchgrass RGB2, the most challenging subset in the test data. See Figure 2 for examples and Table 4 for quantitative results.

**Figure 2:**
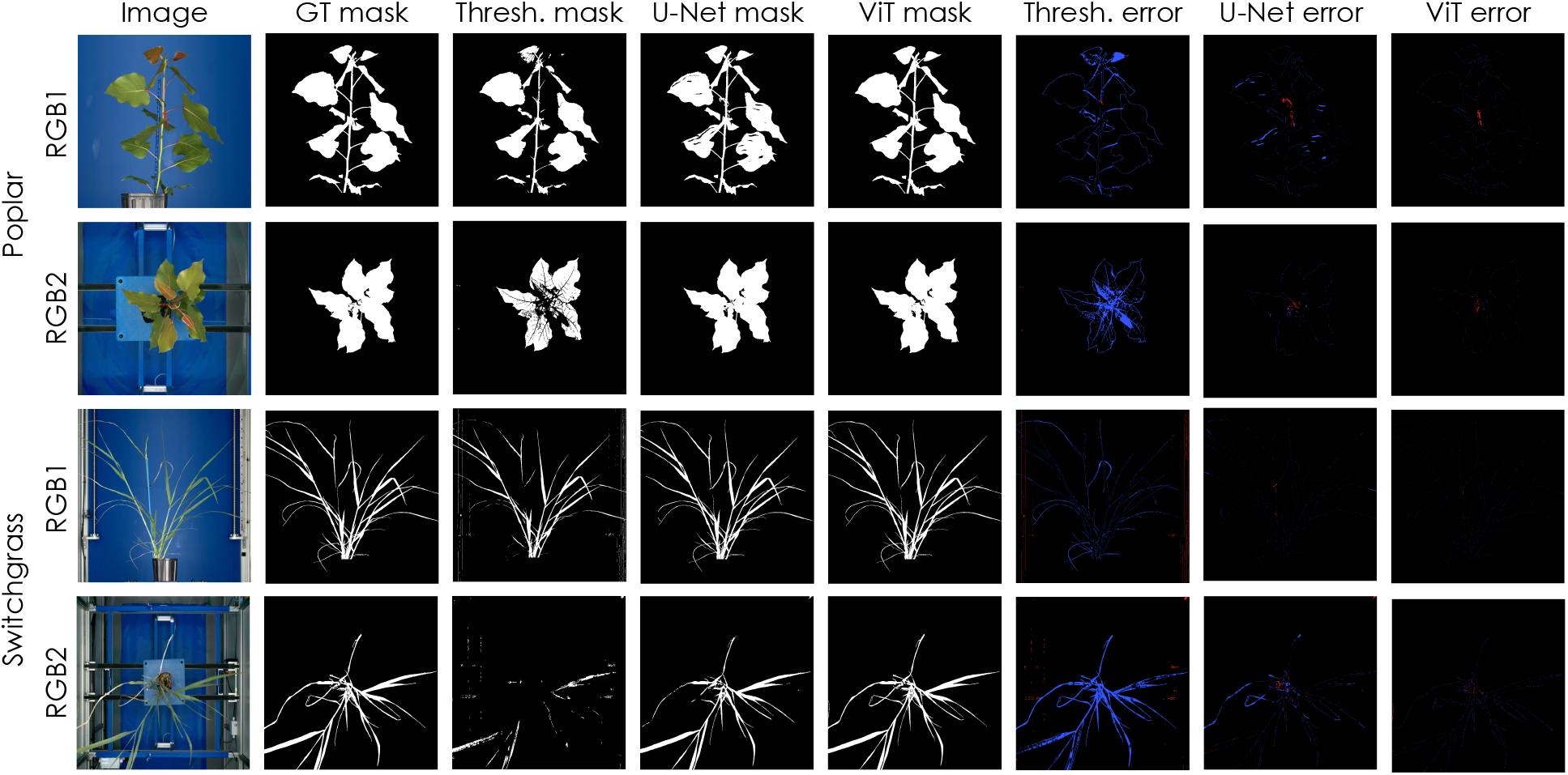
Within-distribution segmentation examples. RGB1 and RGB2 views are shown for each species with the ground-truth (GT) mask, threshold (Thresh.) prediction, U-Net prediction, ViT prediction, and model-specific error maps. Error maps compare each prediction against the ground-truth mask: red indicates false-positive, blue indicates false-negative pixels, and black indicates agreement (true-positive and true-negative pixels). Representative examples were chosen from the test split and cropped to each plant.

**Table 4:** Testing accuracy. Image-level Dice scores (mean ± standard deviation, percentage points) are reported for the threshold-based method and each U-Net and Vision Transformer (ViT) configuration evaluated on the poplar and switchgrass test split. Results are shown overall and separately for each species and imaging modality (RGB1: side-view; RGB2: top-view). Higher Dice values indicate greater overlap between predicted and manually annotated segmentation masks. Best-performing models for each column are shown in bold.

| Strategy | Backbone | Patch | Tile | Overall | Poplar RGB1 | Poplar RGB2 | Switchgrass RGB1 | Switchgrass RGB2 |
| --- | --- | --- | --- | --- | --- | --- | --- | --- |
| Threshold | – | – | – | 58.3 ± 28.5 | 67.3 ± 18.2 | 86.5 ± 9.0 | 55.0 ± 19.4 | 26.1 ± 23.0 |
| U-Net | ResNet-34 | – | 224 | 96.4 ± 4.3 | 97.9 ± 0.8 | 98.7 ± 1.9 | 96.4 ± 1.7 | 92.9 ± 6.8 |
| U-Net | ResNet-34 | – | 448 | 96.6 ± 4.0 | 98.0 ± 0.7 | 98.7 ± 2.0 | 96.5 ± 1.6 | 93.3 ± 6.2 |
| U-Net | ResNet-50 | – | 224 | 96.4 ± 4.4 | 98.0 ± 0.7 | 98.6 ± 2.4 | 96.3 ± 1.7 | 93.0 ± 6.9 |
| U-Net | ResNet-50 | – | 448 | 96.5 ± 4.3 | 98.0 ± 0.7 | 98.7 ± 1.9 | 96.4 ± 1.6 | 93.2 ± 7.0 |
| U-Net | ResNet-101 | – | 224 | 96.1 ± 4.8 | 97.6 ± 1.2 | 98.4 ± 3.4 | 95.9 ± 1.9 | 92.5 ± 7.3 |
| U-Net | ResNet-101 | – | 448 | 96.5 ± 4.0 | 97.9 ± 0.8 | 98.8 ± 1.5 | 96.3 ± 1.6 | 93.3 ± 6.4 |
| U-Net | ResNet-152 | – | 224 | 95.5 ± 5.3 | 97.5 ± 0.8 | 98.2 ± 2.8 | 95.9 ± 1.8 | 90.7 ± 8.2 |
| U-Net | ResNet-152 | – | 448 | 96.3 ± 4.1 | 97.7 ± 0.6 | 98.5 ± 2.4 | 96.3 ± 1.6 | 92.9 ± 6.3 |
| ViT | ViT-Small | 16 | 224 | 96.9 ± 4.0 | 97.9 ± 0.9 | 99.0 ± 0.7 | 96.5 ± 1.6 | 94.3 ± 6.8 |
| ViT | ViT-Small | 16 | 448 | 97.0 ± 3.6 | 97.9 ± 0.8 | 99.1 ± 0.7 | 96.7 ± 1.0 | 94.3 ± 6.1 |
| ViT | ViT-Small | 8 | 224 | **97.3 ± 3.6** | **98.3 ± 0.6** | **99.2 ± 0.5** | 96.8 ± 1.3 | **95.0 ± 6.1** |
| ViT | ViT-Small | 8 | 448 | **97.3 ± 3.6** | **98.3 ± 0.6** | **99.2 ± 0.5** | **96.9 ± 0.9** | 94.9 ± 6.2 |
| ViT | ViT-Base | 16 | 224 | 97.2 ± 3.6 | 98.1 ± 0.7 | 99.1 ± 0.5 | 96.7 ± 1.5 | 94.8 ± 6.2 |
| ViT | ViT-Base | 16 | 448 | 97.2 ± 3.5 | 98.1 ± 0.7 | 99.1 ± 0.6 | 96.8 ± 1.0 | 94.9 ± 6.0 |
| ViT | ViT-Base | 8 | 224 | 97.2 ± 4.0 | **98.3 ± 0.6** | **99.2 ± 0.5** | 96.7 ± 1.6 | 94.9 ± 7.0 |
| ViT | ViT-Base | 8 | 448 | **97.3 ± 3.5** | **98.3 ± 0.6** | **99.2 ± 0.5** | **96.9 ± 1.0** | **95.0 ± 6.0** |

The effects of model capacity and input representation were generally consistent across both architectural families. For the U-Net baselines, increasing the input tile size from 224 to 448 pixels produced consistent improvements in overall Dice, whereas increasing the encoder capacity beyond ResNet-34 did not yield additional gains, slightly reducing accuracy in some cases. ViT performance was also relatively unaffected by backbone size, with both ViT-Small and ViT-Base achieving nearly identical accuracies. Across the evaluated pretrained checkpoints, models using patch size 8 × 8 generally achieved higher Dice than those using 16 × 16. This pattern was also observed within ViT-Base, in which both patch sizes used similar checkpoints, suggesting that finer tokenization preserves small plant structures and boundaries. Similarly, despite differences between the ViT-Small patch-8 and patch-16 model checkpoints, the 8 × 8 patch size consistently outperformed corresponding 16 × 16 variants. Increasing the tile size from 224 to 448 pixels also provided consistent improvements for most ViT configurations, indicating that additional spatial context benefits segmentation while preserving the advantages of pretrained transformer representations. These results suggest that segmentation accuracy depends more on spatial resolution at both the image tile level and the transformer patch level than on increasing encoder capacity alone. Model failures were concentrated along fine structures and object boundaries rather than broad plant/background separation. U-Net errors were most visible as missing thin stems, petioles, and leaf margins, especially in switchgrass RGB2 images, where ViT predictions reduced these errors while preserving the main canopy structure (see Figure 2).

### 3.2 Cross-species generalization

Cross-species evaluation separated the three strategies more clearly. The fixed threshold baseline achieved an overall Dice score of 56.5 ± 32.2, similar to its within-distribution accuracy. The best U-Net achieved 86.2± 23.9 and the best pretrained ViT achieved 95.7 ± 10.1. Relative to thresholding, supervised U-Net provided an improvement of 29.7 percentage points, while pretrained ViT fine-tuning provided a further 9.5-point improvement over the best U-Net.

The threshold baseline was highly context dependent, performing comparatively well on arabidopsis and pennycress, reaching 83.2 and 79.9 Dice, respectively, and exceeding every U-Net variant on those subsets, but fell to 24.2 on eucalyptus and 52.7 on sorghum. This pattern is consistent with a fixed color rule succeeding when plant/background contrast is favorable but failing when that assumption changes. U-Net generalized well on multiple unseen species but remained vulnerable to particular novel top-view scenes and background structures, e.g., new soil backgrounds used in arabidopsis and pennycress experiments. In contrast, the pretrained ViT exceeded 93 mean Dice on every evaluated unseen species, providing the highest accuracy across the tested domain shifts. The species in the generalization dataset differ substantially from the poplar and switchgrass training data in both canopy architecture and leaf morphology, suggesting that pretrained transformer representations capture image features that transfer more effectively across biological domains than supervised CNNs trained from scratch. In addition to improving mean accuracy, pretrained ViTs also consistently reduced the variance in Dice scores, indicating more reliable segmentation across diverse plants, growth stages, and imaging conditions. See Figure 3 for examples and Table 5 for quantitative results.

**Figure 3:**
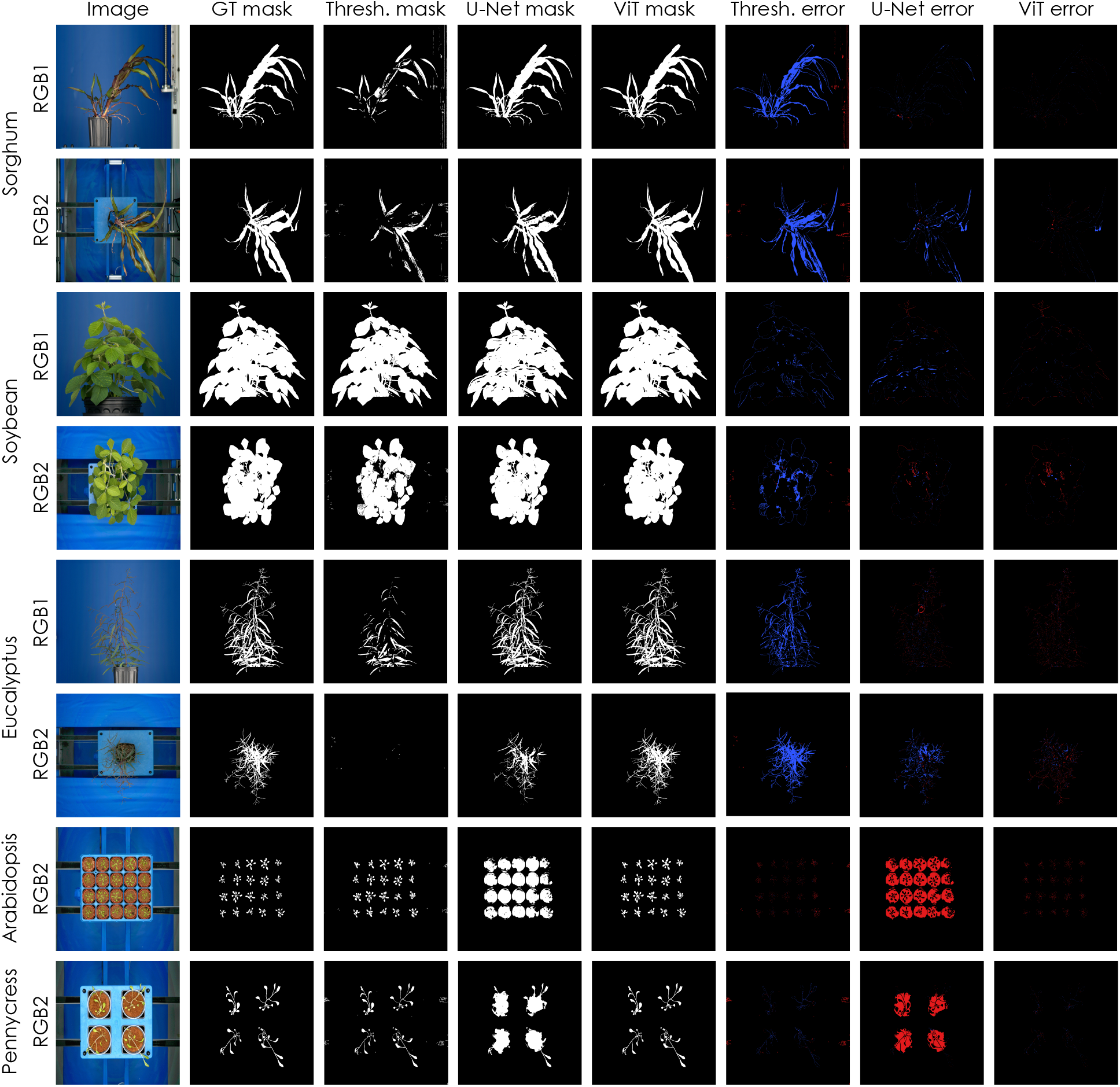
Out-of-distribution segmentation examples. RGB1 and RGB2 views are shown for each species with the ground-truth (GT) mask, threshold (Thresh.) prediction, U-Net prediction, ViT prediction, and model-specific error maps. Error maps compare each prediction against the ground-truth mask: red indicates false-positive pixels, blue indicates false-negative pixels, and black indicates agreement (true-positive and true-negative pixels). Representative examples were chosen from the generalization split and cropped to each plant. Arabidopsis and pennycress included only RGB2 (top-view) images.

**Table 5:** Generalization accuracy. Image-level Dice scores (mean ± standard deviation, percentage points) are reported for the threshold-based method and each U-Net and Vision Transformer (ViT) configuration evaluated on the sorghum, soybean, eucalyptus, arabidopsis, and pennycress generalization split. Results are shown overall and separately for each species. Higher Dice values indicate greater overlap between predicted and manually annotated segmentation masks. Best-performing models for each column are shown in bold.

| Strategy | Backbone | Patch | Tile | Overall | Sorghum | Soybean | Eucalyptus | Arabidopsis | Pennycress |
| --- | --- | --- | --- | --- | --- | --- | --- | --- | --- |
| Threshold | - | - | - | 56.5 ± 32.2 | 52.7 ± 28.9 | 71.3 ± 30.5 | 24.2 ± 20.1 | 83.2 ± 10.2 | 79.9 ± 21.1 |
| U-Net | ResNet-34 | - | 224 | 84.6 ± 26.7 | 88.5 ± 24.4 | 93.4 ± 21.6 | 89.5 ± 9.5 | 60.9 ± 28.8 | 65.6 ± 40.8 |
| U-Net | ResNet-34 | - | 448 | 85.5 ± 26.1 | 89.8 ± 23.7 | 93.9 ± 20.6 | 89.9 ± 9.0 | 62.3 ± 28.6 | 66.4 ± 39.9 |
| U-Net | ResNet-50 | - | 224 | 83.9 ± 27.4 | 88.0 ± 24.9 | 93.3 ± 21.6 | 88.5 ± 12.4 | 58.6 ± 28.9 | 65.3 ± 41.2 |
| U-Net | ResNet-50 | - | 448 | 84.0 ± 27.3 | 88.6 ± 23.8 | 93.4 ± 21.6 | 89.7 ± 10.5 | 54.2 ± 28.9 | 65.0 ± 40.9 |
| U-Net | ResNet-101 | - | 224 | 84.9 ± 25.6 | 87.5 ± 24.7 | 92.6 ± 21.7 | 87.1 ± 13.7 | 71.3 ± 29.6 | 68.7 ± 36.9 |
| U-Net | ResNet-101 | - | 448 | 84.1 ± 27.2 | 88.1 ± 24.5 | 93.2 ± 21.6 | 89.0 ± 12.1 | 58.9 ± 29.2 | 65.4 ± 41.0 |
| U-Net | ResNet-152 | - | 224 | 83.4 ± 25.9 | 86.5 ± 25.1 | 91.9 ± 22.4 | 83.3 ± 16.7 | 70.1 ± 27.6 | 67.7 ± 35.6 |
| U-Net | ResNet-152 | - | 448 | 86.2 ± 23.9 | 89.4 ± 22.4 | 93.6 ± 20.5 | 88.0 ± 12.3 | 69.4 ± 28.8 | 71.5 ± 33.5 |
| ViT | ViT-Small | 16 | 224 | 92.4 ± 17.0 | 91.6 ± 21.8 | 94.2 ± 19.4 | 91.8 ± 9.7 | 91.1 ± 4.8 | 94.0 ± 5.6 |
| ViT | ViT-Small | 16 | 448 | 93.7 ± 13.0 | 94.3 ± 15.9 | 95.7 ± 15.6 | 92.7 ± 3.2 | 89.1 ± 10.2 | 93.6 ± 5.8 |
| ViT | ViT-Small | 8 | 224 | 93.1 ± 16.8 | 90.5 ± 23.7 | 96.5 ± 14.0 | 94.0 ± 7.1 | 92.3 ± 4.5 | 95.4 ± 4.3 |
| ViT | ViT-Small | 8 | 448 | 95.2 ± 12.1 | 95.0 ± 16.8 | **97.0 ± 12.2** | **94.6 ± 2.3** | 93.1 ± 3.1 | 95.5 ± 4.1 |
| ViT | ViT-Base | 16 | 224 | 93.7 ± 14.2 | 92.2 ± 20.2 | 96.9 ± 12.2 | 93.7 ± 2.8 | 92.2 ± 4.3 | 94.8 ± 4.8 |
| ViT | ViT-Base | 16 | 448 | 93.4 ± 14.7 | 91.8 ± 20.5 | 96.4 ± 14.0 | 93.5 ± 2.8 | 92.3 ± 4.4 | 94.8 ± 4.7 |
| ViT | ViT-Base | 8 | 224 | 93.9 ± 16.0 | 92.0 ± 22.9 | **97.0 ± 12.2** | 94.2 ± 7.1 | 93.3 ± 3.7 | 95.6 ± 4.3 |
| ViT | ViT-Base | 8 | 448 | **95.7 ± 10.1** | **96.2 ± 13.0** | **97.0 ± 12.2** | **94.6 ± 2.3** | **93.7 ± 3.3** | **95.7 ± 4.1** |

The trends observed in the test set (see Table 4) became more pronounced for out-of-distribution evaluation in Table 5. Across both U-Net and ViT families, increasing the tile size from 224 to 448 pixels generally improved segmentation performance, suggesting that additional image context becomes increasingly important when segmenting unfamiliar plant morphologies and backgrounds. However, differences in modeling approach remained the dominant factor. Larger ResNet encoders did not consistently improve generalization, with overall Dice varying by less than three percentage points across all CNN configurations. In contrast, ViT models consistently benefited from finer 8 × 8 patch tokenization, with the highest-performing model combining the ViT-Base backbone (the largest ViT variant), 448-pixel input tiles (the largest tile size), and 8-pixel patches (the smallest patch size). This configuration achieved the best overall performance as well as the highest Dice on sorghum, arabidopsis, and pennycress, while tying for the highest performance on soybean and eucalyptus. These results indicate that preserving finer spatial detail and increasing image context provide complementary benefits for pretrained transformer-based segmentation, whereas simply increasing CNN encoder capacity is insufficient to overcome domain shifts encountered in realistic plant phenotyping experiments. Generalization accuracy revealed larger differences between model families, where U-Net often over-segmented non-plant tray or pot regions, e.g., in arabidopsis and pennycress, while missing fine leaf structures in eucalyptus. ViT errors were generally sparser and localized to thin leaves, occluded boundaries, and small plant structures, indicating that most of the generalization gains in accuracy came from suppressing false positives on unfamiliar backgrounds while retaining finer plant morphology. The threshold-based method was largely dependent on plant pigmentation, with higher segmentation accuracy for soybean, arabidopsis, and pennycress, and significant error rates for sorghum and eucalyptus. See Supplementary Table 1 for the full generalization metrics by species and modality, which further illustrate the context sensitivity of fixed thresholding. Threshold-based Dice ranged from 9.7 on eucalyptus RGB2 to 87.0 on soybean RGB2, whereas the best pretrained ViTs remained above 91 Dice for every species–modality combination.

### 3.3 Tile aggregation accuracy and efficiency

Hann-windowed overlapping inference consistently improved segmentation accuracy relative to classical non-overlapping tiling across all segmentation models evaluated (Supplementary Table 2). Averaged across all 16 model configurations, Hann reconstruction increased image-level Dice by 3.23%. Improvements were observed for both U-Net and ViT model families, though the magnitude varied substantially across backbone and tile-size combinations. The largest overall improvement was obtained by the smallest U-Net (ResNet-34 backbone) with 224 × 224 tiles with a gain of 13.2%. Among the ViT models, gains were consistently positive but smaller, ranging from +1.2% to +3.0%, with the largest improvement achieved by the ViT-Base model using patch-16 and 448 × 448 tiles. Accuracy gains were generally larger for the higher-resolution RGB1 (side-view) modality than for RGB2 (top-view). For example, U-Net with ResNet-34 and 224 × 224 tiles improved by +28.8% on RGB1 images but only +2.8% on RGB2 images. Overall, these results demonstrate that Hann-windowed reconstruction consistently improves segmentation accuracy across architectures, with the greatest benefits observed for larger, high-resolution images.

The accuracy gains by the overlapping Hann-window method were accompanied by larger inference times because overlapping inference evaluates additional image tiles and blends their probability maps (Supplementary Table 3). Averaged across all model configurations, Hann reconstruction increased inference time from 1.36 to 2.65 s/image, corresponding to an average increase of 1.28 s while improving Dice by 3.23 percentage points. The overhead was greater for RGB1 side-view images, where inference time increased from 2.04 to 4.03 s/image (+2.00 s), than for RGB2 top-view images, where inference time increased from 0.91 to 1.72 s/image (+0.81 s). Among all configurations, U-Net with ResNet-50 and 224 × 224 tiles was the fastest under Hann inference (1.74 ± 0.70 s/image), where the best performing ViT model (ViT-Base with 8-pixel patches and 448 × 448 tiles) required the longest inference time (5.26 ± 2.10 s/image). Thus, Hann-windowed inference approximately doubled inference time while providing a consistent improvement in segmentation accuracy across all evaluated architectures.

## 4 Discussion

This study evaluated three practical plant-segmentation strategies that use progressively richer visual priors: (i) fixed color thresholding, (ii) fully supervised U-Net training, and (iii) fine-tuning pretrained ViTs. The largest improvement on the held-out poplar and switchgrass test set came from replacing the fixed rule with learned segmentation, increasing overall mean Dice from 58.3 to 96.6. The distinction between the two learned strategies became much larger under cross-species evaluation, where the best U-Net reached 86.2 and the best pretrained ViT reached 95.7 mean Dice.

The threshold method is an informative baseline since it requires no labeled training data, is transparent, and performed well on several top-view datasets. In particular, thresholding exceeded every evaluated U-Net on arabidopsis and pennycress. Its overall performance, however, was highly variable and failed for eucalyptus, switchgrass RGB2, and multiple sorghum conditions. These differences capture the trade-offs of using fixed color rules, where as the plant tissue and background match the fixed rule, thresholding can be effective, but performance may not transfer when plant pigmentation, soil type, or background artifacts change. Thresholding may therefore remain appropriate for stable and well-characterized imaging and experimental configurations, while requiring retuning or replacement as experimental conditions change.

Supervised U-Nets produced the largest absolute gain over thresholding and achieved high accuracy on species represented during training. Across the full generalization set, these models also substantially outperformed thresholding, demonstrating the value of learning spatial and contextual features from manual masks rather than relying on fixed color rules. However, they generalized less consistently, e.g., failing for arabidopsis and pennycress experiments, in which simple thresholding remained suitable while U-Net frequently classified unfamiliar backgrounds as plants. Competitive within-distribution performance therefore did not guarantee generalization to novel experimental conditions, and supervised learning may have encoded visual representations that were less stable than the threshold-based rules for particular cases.

Fine-tuned pretrained ViTs combined the high within-distribution accuracy of learned segmentation with the most consistent performance to unseen species and experimental conditions. The advantage of pretrained ViTs was not primarily the minor improvement over U-Net on held-out poplar and switchgrass, where both learned models were highly accurate. Rather, it was the substantial improvement on the generalization set and the absence of subsets of failures observed for both thresholding and U-Net. The best ViT exceeded 93 mean Dice on all unseen species, while retaining thin leaves and plant boundaries and suppressing false positives from unfamiliar backgrounds. This pattern suggests pretrained visual features may provide information that transfers more broadly than representations learned only from the APPL training masks.

The learned model comparisons also show that spatial context and feature resolution are more important than model size/capacity alone. Larger ResNet backbones did not consistently improve either test or generalization accuracy, and several deeper U-Net configurations performed similarly or worse than the smaller ResNet-34 variants (Tables 4 and 5). This occurred despite the substantial increase in parameter count across the ResNet encoders (Table 2). ViT-Small and ViT-Base likewise produced nearly identical results on the within-distribution test set, though ViT-Base achieved the highest overall generalization accuracy when paired with the largest input tiles and smallest patch size. Across both model families, 448 × 448-pixel tiles generally outperformed 224 × 224-pixel tiles, suggesting that broader image context helps distinguish plant structures from visually similar background objects. Within the transformer models, 8 × 8 patches generally outperformed 16 × 16 patches, indicating that finer tokenization better preserves complex plant structures. The best-performing configuration therefore combined two complementary forms of spatial information: a large tile size provided sufficient contextual information to interpret the surrounding scene, while small patch size retained the detail required for precise segmentation. These findings indicate that simply increasing model size is unlikely to resolve domain shifts in phenotyping unless the input representation also preserves the spatial scales relevant to plant morphology.

The performance of the ViT models is notable given that fine-tuning used only the 1,500 images in the training split, which was divided between the two APPL RGB modalities and plant species (Table 1). Large manually segmented datasets are prohibitively expensive to produce, particularly for full-resolution phenotyping images containing complex canopy architectures, fine plant structures, and multiple plants per image [5]. Large-scale, self-supervised pretraining may provide a method for reducing this annotation bottleneck by providing generalizable visual representations learned from massive image datasets before task-specific fine-tuning [7, 11, 12]. The consistent ViT performance across the unseen species in Table 5 suggests that these pretrained features include transferable information about colors, shapes, textures, and object-level context that was not learned as reliably by a CNN baseline trained from scratch. However, the present comparison does not isolate the effect of the ViT architecture itself from the effect of pretraining, because ViT encoders were pretrained whereas the U-Net encoders were randomly initialized, as described in Section 2.2. The results should therefore be interpreted as a comparison between two practical training strategies (fine-tuning pretrained ViTs and supervised training of U-Nets described in Section 2.3) rather than evidence that self-attention in transformers alone explains the improvements in segmentation accuracy. The results suggest that fine-tuning pretrained ViT encoders is a practical strategy for controlled-environment plant segmentation, potentially reducing annotation requirements while maintaining strong performance across biologically diverse species and imaging conditions.

Hann-weighted tile overlapping further provided an architecture-independent improvement in segmentation quality. Across all model configurations, overlapping reconstruction increased image-level Dice by an average of 3.23 percentage points relative to non-overlapping tiling (Supplementary Table 2). The benefit was greatest for models using smaller tiles and for the higher-resolution side-view RGB1 images, where a larger number of independently predicted tile boundaries were introduced during full-image reconstruction. These results support the interpretation that predictions near tile edges are less reliable because the model has access to less surrounding context than it does near tile centers (Section 2.4). Non-overlapping reconstruction gives these edge predictions the same weight as predictions near tile centers, producing discontinuities and repeated artifacts along the tiling grid. In contrast, Hann weighting emphasizes the central region of each tile and smoothly combines probabilities from overlapping predictions before assigning the final plant/background classification (Supplementary Figure 2). Importantly, this improvement required no model retraining, changes to model architecture, or morphological post-processing.

The increased accuracy obtained through model family, larger tiles, and overlapping inference is compared with computational throughput. Hann-weighted reconstruction approximately doubled average inference time from 1.36 to 2.65 seconds per image while increasing Dice by an average of 3.23 percentage points (Supplementary Tables 2 and 3). The most accurate configuration, ViT-Base with 8 × 8 patches and 448 × 448-pixel tiles, achieved the highest overall generalization Dice in Table 5 but required 5.26 ± 2.10 seconds per image under Hann-weighted inference. This additional cost results primarily from evaluating overlapping tiles and, for ViT-Base, processing a larger ViT encoder with finer patch tokenization. The appropriate configuration therefore depends on the downstream application. High-accuracy reconstruction may be preferable for final trait extraction and experiments in which segmentation errors could bias biological conclusions. Smaller ViT models, smaller tiles, or non-overlapping reconstruction may remain useful for rapid quality control, preliminary visualization, or real-time applications with latency constraints. Although the highest-performing configurations increase per-image inference latency, the workload is inherently parallelizable since each image is processed independently. Therefore, overall throughput can be scaled across multiple GPUs, making these models practical for large high-throughput phenotyping experiments despite their higher computational cost.

Several limitations define the scope of these conclusions. For example, all images considered in this work were collected within APPL, meaning that even though the generalization dataset represents new species, plant morphology, growth stages, tray types, and soil backgrounds beyond the training data, it does not evaluate generalization across phenotyping facilities, camera systems, or lighting conditions. Further, the models were evaluated only on RGB imagery and binary plant/background segmentation. It is unclear whether these results extend to additional APPL imaging modalities, including hyperspectral, chlorophyll fluorescence, thermal, and multispectral (Section 1). Similarly, the manually curated generalization dataset included diverse images, but it does not represent every possible APPL experiment or domain shift. Each learned model configuration was also trained only once, meaning that small differences among closely performing configurations should be interpreted descriptively and may not quantify variability across random initializations or training runs. The image-level Dice metric described in Section 2.4 measures overall predicted/ground truth overlap but does not directly quantify the resulting error in plant phenotypes like projected leaf area, plant height, growth rate, or stress response. This quantification would require additional manual phenotype measurements that are beyond the scope of this work. Finally, no object filtering, hole filling, or other segmentation post-processing was applied during evaluation of the learned models in order to enable a controlled comparison of the raw model predictions in Figures 2 and 3, but more sophisticated phenotyping pipelines may obtain additional improvements from carefully validated post-processing methods, similar to the APPL threshold algorithm described in Section 2.2.

Future work will evaluate how these models transfer across facilities, modalities, species, and experimental conditions that differ more substantially from the APPL data described in Section 2.1. Comparisons among pretrained ViT encoders, including general-purpose segmentation foundation models like the Segment Anything Model (SAM) family [36, 37, 38] and modern self-supervised training paradigms like Joint-Embedding Predictive Architectures (JEPA) [39, 40] would help distinguish the contributions of architecture, pretraining objective, and pretraining data. Few-shot learning strategies could further reduce the annotation requirements established by the 1,500-image training split [5]. Each plant in APPL is imaged over time, incorporating temporal information may therefore also improve consistency across time-resolved plant images, where unrealistic changes in plant area or shape could be detected using neighboring observations. Extending the framework to the additional APPL modalities may provide more robust segmentation when RGB contrast is low. Model evaluation will also require moving beyond computer vision metrics like the Dice coefficient in order to quantify how segmentation accuracy affects downstream prediction of plant size, morphology, color, and growth. Finally, model distillation, adaptive tiling, and optimized inference could maintain within-distribution and out-of-distribution segmentation accuracy while reducing the computational costs required by GPU-based inference. Together, these developments could support reusable segmentation models that operate across experiments with limited additional annotation, shifting controlled-environment phenotyping toward scalable and automated trait extraction, reducing the time it takes to go from experiment to insight and thereby accelerating scientific discovery in plant biology.

## 5 Conclusions

This work showed that fixed thresholding can be effective in stable, color-separable imaging conditions but is unreliable across heterogeneous species and backgrounds. Supervised U-Net provides a large improvement over fixed rules and excellent accuracy in the trained domain, but remains sensitive to changes in experimental conditions. Fine-tuning pretrained ViTs produced the most consistently high performance across the evaluated species, views, tray types, and plant growth stages. This broad improvement is the primary reason to favor pretrained visual representations for reusable controlled-environment phenotyping pipelines.

## 6 Acknowledgments

This material is based upon work at the Center for Bioenergy Innovation (CBI) supported by the U.S. Department of Energy, Office of Science, Biological and Environmental Research Program under Contract Number ERKP886. This research used resources at the Advanced Plant Phenotyping Laboratory (APPL), a shared-use facility operated by the Oak Ridge National Laboratory. This research was partially sponsored by the Laboratory Directed Research and Development Program of Oak Ridge National Laboratory, managed by UT-Battelle, LLC, for the US Department of Energy. Notice: This manuscript has been authored by UT-Battelle, LLC, under contract DE-AC05-00OR22725 with the US Department of Energy (DOE). The US government retains and the publisher, by accepting the article for publication, acknowledges that the US government retains a nonexclusive, paid-up, irrevocable, worldwide license to publish or reproduce the published form of this manuscript, or allow others to do so, for US government purposes. DOE will provide public access to these results of federally sponsored research in accordance with the DOE Public Access Plan (https://www.energy.gov/doe-public-access-plan).

## 8 AI use disclosure

AI tools, including GitHub Copilot and OpenAI ChatGPT/Codex, were used as pair-programming and drafting aids during this work. They supported code inspection, debugging, code edits, documentation, manuscript editing, and text refinement. AI was not used as an autonomous decision-maker or as a source of unverified scientific claims. The research design, code, analyses, results, and conclusions have been reviewed and edited for content and accuracy by the authors, who accept full responsibility for the work.

## 9 Author contributions (CRediT)

J.M., A.S., L.Y., and J.L. conceptualized the study, designed the methodology, and conducted the investigation. L.Y. and J.L. procured the data. J.M., A.S., L.Y., and J.L. curated the data. J.M. and J.L. implemented the software and performed the formal analysis. A.T. and X.W. provided valuable feedback during the development process. J.M. and J.L. wrote the original draft of the manuscript, and all authors contributed to review and editing. L.Y. and J.L. supervised and administered the project, and acquired funding.

## 10 Data availability

Upon publication, the source code will be publicly available on GitHub, with a versioned archival copy preserved in Zenodo. The images, ground-truth segmentation masks, and trained model checkpoints will be publicly available through Zenodo and/or the U.S. Department of Energy Office of Scientific and Technical Information (OSTI).

