## Supplementary for "Vision Transformers Enable Advanced Plant Phenotyping in Controlled Environments"


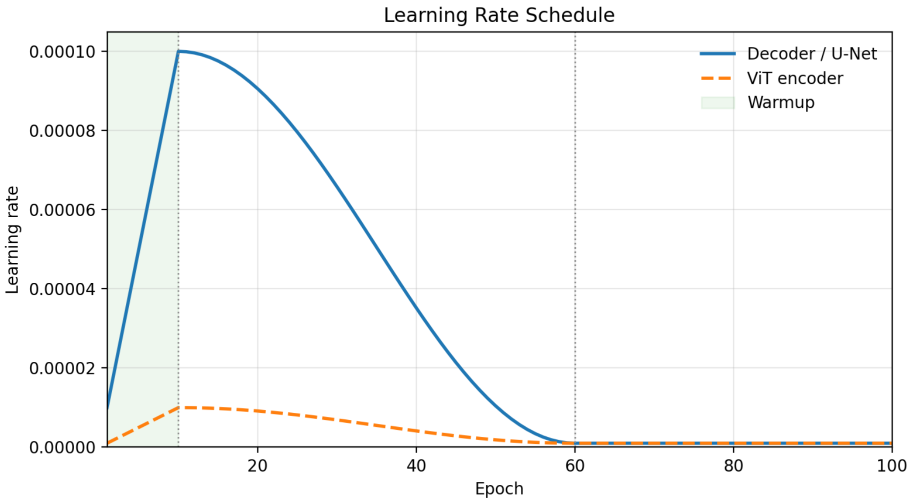


Figure 1: **Learning rate schedule.** All models were trained for 100 epochs using a learning rate schedule consisting of a 10-epoch linear warmup, a 50-epoch cosine decay, and a final constant learning rate through epoch 100. U-Net models used a single learning rate schedule with a maximum learning rate of $1 \times 10^{-4}$ and a minimum learning rate of $1 \times 10^{-6}$. For Vision Transformer (ViT) models, the same schedule was applied to both the pretrained encoder and randomly initialized decoder, with the maximum encoder learning rate scaled by a factor of 0.1 relative to the decoder throughout training to enable more conservative fine-tuning of pretrained weights.


Notice: This manuscript has been authored by UT-Battelle, LLC, under contract DE-AC05-00OR22725 with the US Department of Energy (DOE). The US government retains and the publisher, by accepting the article for publication, acknowledges that the US government retains a nonexclusive, paid-up, irrevocable, worldwide license to publish or reproduce the published form of this manuscript, or allow others to do so, for US government purposes. DOE will provide public access to these results of federally sponsored research in accordance with the DOE Public Access Plan (`https://www.energy.gov/doe-public-access-plan`).

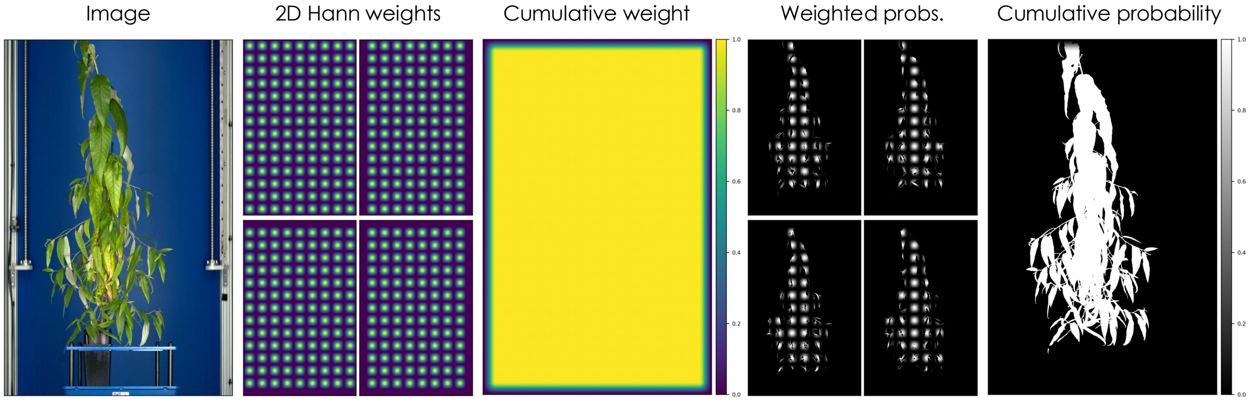


Figure 2: **Inference algorithm.** During inference, each full-resolution image was divided into overlapping tiles with a 50% stride and independently segmented by the model. A two-dimensional (2D) Hann window was applied to each tile probability map before reconstruction, assigning greater weight to predictions near tile centers and lower weight to predictions near tile boundaries. The weighted probabilities from all overlapping tiles were accumulated and normalized by the cumulative Hann weights to produce a full-image probability map. The final segmentation mask was obtained by assigning each pixel to the class with the highest normalized probability, reducing visible stitching artifacts at tile boundaries.

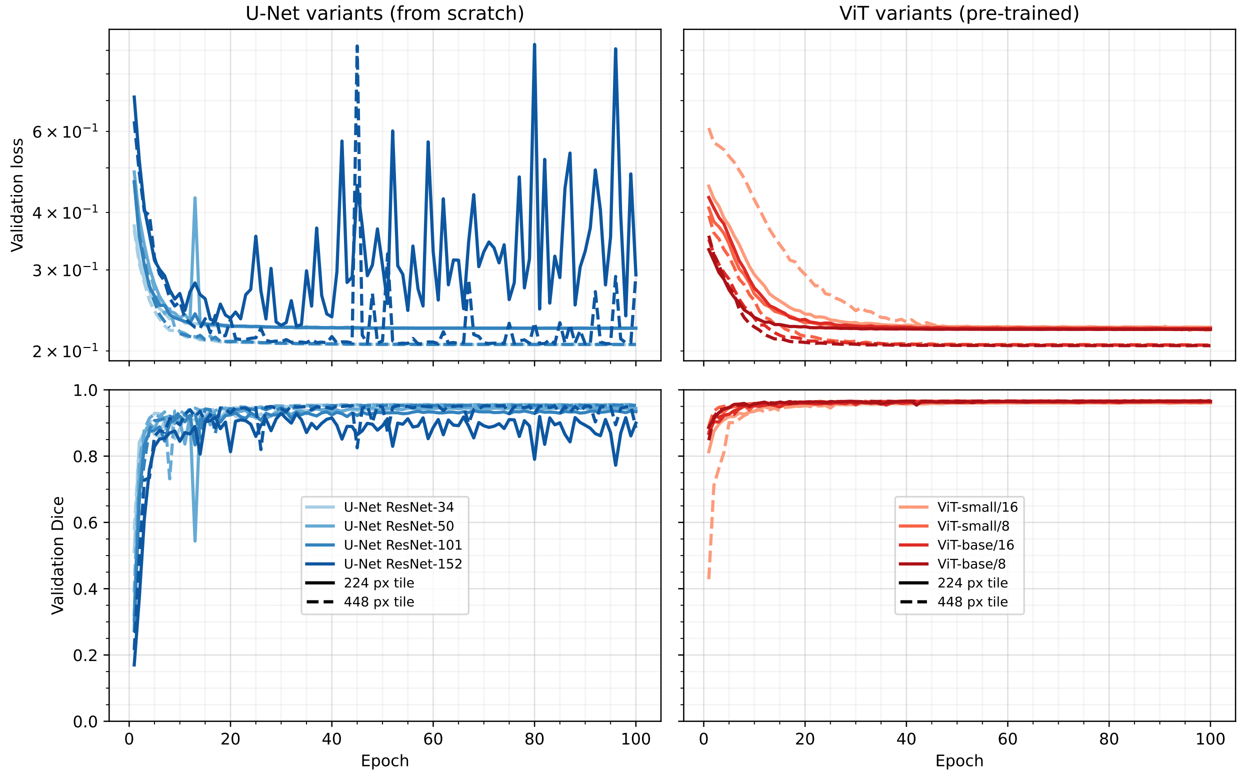


Figure 3: **Model convergence.** Validation loss and validation Dice are shown across training epochs for U-Net variants trained from scratch and pre-trained Vision Transformer (ViT) variants. U-Net models use ResNet-34, ResNet-50, ResNet-101, and ResNet-152 encoders; ViT models use small and base backbones with patch sizes of 8 or 16. Colors distinguish model variants, and line styles distinguish tile sizes of 224 and 448 pixels. Lower validation loss and higher validation Dice indicate better performance.

Table 1: **Generalization accuracy by species and modality.** Image-level Dice scores (mean ± standard deviation, percentage points) are reported for the threshold-based method and each U-Net and Vision Transformer (ViT) configuration evaluated on the generalization split. Supplemental results are shown for every observed species-modality pair. Higher Dice values indicate greater overlap between predicted and manually annotated segmentation masks. Best-performing models for each column are shown in bold.

| Backbone | Patch | Tile | Overall | Sorghum RGB1 | Sorghum RGB1 Rhizo | Sorghum RGB2 | Sorghum RGB2 Rhizo |
|---|---|---|---|---|---|---|---|
| Threshold | - | - | $56.5 \pm 32.2$ | $53.7 \pm 24.2$ | $33.7 \pm 29.8$ | $72.2 \pm 17.3$ | $51.1 \pm 28.9$ |
| ResNet-34 | - | 224 | $84.6 \pm 26.7$ | $97.5 \pm 1.5$ | $75.9 \pm 32.7$ | $97.0 \pm 2.2$ | $83.6 \pm 31.3$ |
| ResNet-34 | - | 448 | $85.5 \pm 26.1$ | $97.7 \pm 1.5$ | $80.6 \pm 32.2$ | $97.1 \pm 2.1$ | $83.7 \pm 31.4$ |
| ResNet-50 | - | 224 | $83.9 \pm 27.4$ | $97.6 \pm 1.6$ | $73.5 \pm 33.2$ | $97.0 \pm 2.2$ | $83.8 \pm 31.4$ |
| ResNet-50 | - | 448 | $84.0 \pm 27.3$ | $97.7 \pm 1.6$ | $71.7 \pm 33.6$ | $97.3 \pm 2.1$ | $87.8 \pm 26.4$ |
| ResNet-101 | - | 224 | $84.9 \pm 25.6$ | $96.3 \pm 3.3$ | $73.5 \pm 32.9$ | $96.6 \pm 2.5$ | $83.6 \pm 31.3$ |
| ResNet-101 | - | 448 | $84.1 \pm 27.2$ | $97.3 \pm 2.3$ | $74.7 \pm 32.7$ | $97.1 \pm 2.1$ | $83.5 \pm 31.3$ |
| ResNet-152 | - | 224 | $83.4 \pm 25.9$ | $96.7 \pm 2.6$ | $71.3 \pm 33.4$ | $94.8 \pm 4.3$ | $83.1 \pm 31.2$ |
| ResNet-152 | - | 448 | $86.2 \pm 23.9$ | $97.6 \pm 1.5$ | $74.9 \pm 32.7$ | $96.3 \pm 2.8$ | $88.8 \pm 24.8$ |
| ViT-Small | 16 | 224 | $92.4 \pm 17.0$ | $97.7 \pm 1.5$ | $81.8 \pm 31.9$ | $98.4 \pm 0.7$ | $88.7 \pm 26.5$ |
| ViT-Small | 16 | 448 | $93.7 \pm 13.0$ | $97.6 \pm 1.5$ | $84.8 \pm 29.7$ | $98.4 \pm 1.1$ | $\mathbf{96.6 \pm 3.2}$ |
| ViT-Small | 8 | 224 | $93.1 \pm 16.8$ | $\mathbf{98.0 \pm 1.4}$ | $82.2 \pm 31.9$ | $98.6 \pm 0.8$ | $83.4 \pm 31.5$ |
| ViT-Small | 8 | 448 | $95.2 \pm 12.1$ | $\mathbf{98.0 \pm 1.4}$ | $94.2 \pm 19.4$ | $98.6 \pm 0.6$ | $89.2 \pm 26.6$ |
| ViT-Base | 16 | 224 | $93.7 \pm 14.2$ | $97.8 \pm 1.4$ | $79.6 \pm 32.1$ | $98.6 \pm 0.7$ | $92.8 \pm 19.3$ |
| ViT-Base | 16 | 448 | $93.4 \pm 14.7$ | $97.8 \pm 1.5$ | $78.1 \pm 32.5$ | $98.6 \pm 0.6$ | $92.8 \pm 19.3$ |
| ViT-Base | 8 | 224 | $93.9 \pm 16.0$ | $\mathbf{98.0 \pm 1.4}$ | $86.4 \pm 30.7$ | $\mathbf{98.7 \pm 0.6}$ | $84.9 \pm 31.6$ |
| ViT-Base | 8 | 448 | $\mathbf{95.7 \pm 10.1}$ | $97.9 \pm 1.3$ | $\mathbf{96.2 \pm 13.9}$ | $\mathbf{98.7 \pm 0.6}$ | $91.9 \pm 21.5$ |

| Backbone | Patch | Tile | Soybean RGB1 | Soybean RGB2 | Eucalyptus RGB1 | Eucalyptus RGB2 | Arabidopsis RGB2 | Pennycress RGB2 |
|---|---|---|---|---|---|---|---|---|
| Threshold | - | - | $55.5 \pm 32.2$ | $87.0 \pm 18.4$ | $38.7 \pm 17.2$ | $9.7 \pm 9.5$ | $83.2 \pm 10.2$ | $79.9 \pm 21.1$ |
| ResNet-34 | - | 224 | $91.7 \pm 25.3$ | $95.0 \pm 17.1$ | $93.4 \pm 10.0$ | $85.7 \pm 7.2$ | $60.9 \pm 28.8$ | $65.6 \pm 40.8$ |
| ResNet-34 | - | 448 | $92.7 \pm 23.6$ | $95.0 \pm 17.1$ | $93.5 \pm 9.9$ | $86.4 \pm 6.4$ | $62.3 \pm 28.6$ | $66.4 \pm 39.9$ |
| ResNet-50 | - | 224 | $91.7 \pm 25.3$ | $95.0 \pm 17.2$ | $91.3 \pm 15.4$ | $85.6 \pm 7.5$ | $58.6 \pm 28.9$ | $65.3 \pm 41.2$ |
| ResNet-50 | - | 448 | $91.7 \pm 25.3$ | $95.1 \pm 17.0$ | $92.0 \pm 13.0$ | $87.4 \pm 6.4$ | $54.2 \pm 28.9$ | $65.0 \pm 40.9$ |
| ResNet-101 | - | 224 | $90.2 \pm 25.3$ | $94.9 \pm 17.2$ | $90.3 \pm 17.1$ | $83.8 \pm 8.1$ | $71.3 \pm 29.6$ | $68.7 \pm 36.9$ |
| ResNet-101 | - | 448 | $91.2 \pm 25.3$ | $95.3 \pm 17.0$ | $91.2 \pm 15.7$ | $86.7 \pm 6.2$ | $58.9 \pm 29.2$ | $65.4 \pm 41.0$ |
| ResNet-152 | - | 224 | $91.1 \pm 25.2$ | $92.7 \pm 19.2$ | $90.0 \pm 17.4$ | $76.5 \pm 13.1$ | $70.1 \pm 27.6$ | $67.7 \pm 35.6$ |
| ResNet-152 | - | 448 | $92.2 \pm 23.4$ | $94.9 \pm 17.1$ | $91.3 \pm 14.9$ | $84.8 \pm 7.8$ | $69.4 \pm 28.8$ | $71.5 \pm 33.5$ |
| ViT-Small | 16 | 224 | $93.4 \pm 21.6$ | $95.0 \pm 17.1$ | $91.7 \pm 13.5$ | $91.9 \pm 2.8$ | $91.1 \pm 4.8$ | $94.0 \pm 5.6$ |
| ViT-Small | 16 | 448 | $96.4 \pm 13.9$ | $95.0 \pm 17.1$ | $93.6 \pm 3.3$ | $91.8 \pm 2.8$ | $89.1 \pm 10.2$ | $93.6 \pm 5.8$ |
| ViT-Small | 8 | 224 | $97.7 \pm 9.9$ | $95.3 \pm 17.1$ | $94.6 \pm 9.7$ | $93.4 \pm 2.4$ | $92.3 \pm 4.5$ | $95.4 \pm 4.3$ |
| ViT-Small | 8 | 448 | $\mathbf{98.7 \pm 1.1}$ | $95.3 \pm 17.1$ | $\mathbf{95.6 \pm 2.0}$ | $93.6 \pm 2.1$ | $93.1 \pm 3.1$ | $95.5 \pm 4.1$ |
| ViT-Base | 16 | 224 | $98.5 \pm 1.6$ | $95.2 \pm 17.1$ | $94.5 \pm 2.8$ | $92.8 \pm 2.4$ | $92.2 \pm 4.3$ | $94.8 \pm 4.8$ |
| ViT-Base | 16 | 448 | $97.5 \pm 10.0$ | $95.2 \pm 17.1$ | $94.4 \pm 2.7$ | $92.7 \pm 2.5$ | $92.3 \pm 4.4$ | $94.8 \pm 4.7$ |
| ViT-Base | 8 | 224 | $\mathbf{98.7 \pm 1.2}$ | $95.3 \pm 17.1$ | $94.5 \pm 9.8$ | $\mathbf{94.0 \pm 2.0}$ | $93.3 \pm 3.7$ | $95.6 \pm 4.3$ |
| ViT-Base | 8 | 448 | $\mathbf{98.7 \pm 1.2}$ | $\mathbf{95.4 \pm 17.0}$ | $95.4 \pm 2.1$ | $93.8 \pm 2.1$ | $\mathbf{93.7 \pm 3.3}$ | $\mathbf{95.7 \pm 4.1}$ |

Table 2: **Inference accuracy.** Image-level Dice-score gains (Hann minus classical inference, mean ± standard deviation, percentage points) are reported for each U-Net and Vision Transformer (ViT) configuration evaluated on the generalization split. RGB1 includes RGB1 and RGB1-rhizo images; RGB2 includes RGB2 and RGB2-rhizo images. Higher values indicate larger gains from Hann-windowed inference. Best gains for each column are shown in bold.

| Backbone | Patch | Tile | Overall | RGB1 | RGB2 |
|---|---|---|---|---|---|
| ResNet-34 | - | 224 | $+\mathbf{13.2} \pm \mathbf{20.4}$ | $+\mathbf{28.8} \pm \mathbf{24.1}$ | $+2.8 \pm 6.1$ |
| ResNet-34 | - | 448 | $+1.0 \pm 5.3$ | $+1.7 \pm 8.1$ | $+0.5 \pm 1.6$ |
| ResNet-50 | - | 224 | $+1.1 \pm 3.6$ | $+0.9 \pm 3.2$ | $+1.2 \pm 3.8$ |
| ResNet-50 | - | 448 | $+3.9 \pm 9.8$ | $+5.2 \pm 9.2$ | $+3.0 \pm 10.2$ |
| ResNet-101 | - | 224 | $+4.2 \pm 7.6$ | $+5.9 \pm 8.7$ | $+3.1 \pm 6.5$ |
| ResNet-101 | - | 448 | $+1.2 \pm 3.7$ | $+1.0 \pm 2.4$ | $+1.4 \pm 4.4$ |
| ResNet-152 | - | 224 | $+2.1 \pm 3.9$ | $+1.9 \pm 3.2$ | $+2.3 \pm 4.3$ |
| ResNet-152 | - | 448 | $+9.2 \pm 16.7$ | $+17.8 \pm 20.4$ | $+\mathbf{3.5} \pm \mathbf{10.3}$ |
| ViT-Small | 16 | 224 | $+1.2 \pm 8.0$ | $+1.1 \pm 7.7$ | $+1.3 \pm 8.2$ |
| ViT-Small | 16 | 448 | $+1.7 \pm 12.7$ | $+0.4 \pm 10.2$ | $+2.5 \pm 14.0$ |
| ViT-Small | 8 | 224 | $+1.3 \pm 8.6$ | $+2.1 \pm 13.2$ | $+0.8 \pm 2.3$ |
| ViT-Small | 8 | 448 | $+2.1 \pm 12.9$ | $+3.4 \pm 17.7$ | $+1.2 \pm 8.2$ |
| ViT-Base | 16 | 224 | $+2.6 \pm 13.3$ | $+3.9 \pm 15.6$ | $+1.7 \pm 11.4$ |
| ViT-Base | 16 | 448 | $+3.0 \pm 13.1$ | $+4.3 \pm 15.0$ | $+2.0 \pm 11.6$ |
| ViT-Base | 8 | 224 | $+1.4 \pm 9.8$ | $+3.1 \pm 15.3$ | $+0.4 \pm 0.6$ |
| ViT-Base | 8 | 448 | $+2.5 \pm 15.3$ | $+4.4 \pm 20.2$ | $+1.3 \pm 10.7$ |

Table 3: **Inference time.** Per-image inference time (mean ± standard deviation, seconds) is reported for each U-Net and Vision Transformer (ViT) configuration evaluated on the generalization split. RGB1 includes RGB1 and RGB1-rhizo images; RGB2 includes RGB2 and RGB2-rhizo images. Lower values indicate faster inference. Fastest models for each column are shown in bold.

| Backbone | Patch | Tile | Overall | | RGB1 | | RGB2 | |
|---|---|---|---|---|---|---|---|---|
| | | | Hann | Classical | Hann | Classical | Hann | Classical |
| ResNet-34 | - | 224 | $1.77 \pm 0.79$ | $1.13 \pm 0.63$ | $2.73 \pm 0.19$ | $1.68 \pm 0.71$ | $\mathbf{1.13 \pm 0.04}$ | $\mathbf{0.76 \pm 0.03}$ |
| ResNet-34 | - | 448 | $2.06 \pm 0.83$ | $1.22 \pm 0.65$ | $3.08 \pm 0.04$ | $1.86 \pm 0.60$ | $1.38 \pm 0.03$ | $0.79 \pm 0.03$ |
| ResNet-50 | - | 224 | $\mathbf{1.74 \pm 0.70}$ | $1.14 \pm 0.74$ | $\mathbf{2.60 \pm 0.04}$ | $1.72 \pm 0.90$ | $1.16 \pm 0.04$ | $\mathbf{0.76 \pm 0.03}$ |
| ResNet-50 | - | 448 | $2.03 \pm 0.83$ | $1.19 \pm 0.74$ | $3.05 \pm 0.04$ | $1.75 \pm 0.92$ | $1.35 \pm 0.03$ | $0.81 \pm 0.02$ |
| ResNet-101 | - | 224 | $2.04 \pm 0.97$ | $\mathbf{1.11 \pm 0.77}$ | $3.22 \pm 0.08$ | $\mathbf{1.63 \pm 1.01}$ | $1.25 \pm 0.02$ | $\mathbf{0.76 \pm 0.01}$ |
| ResNet-101 | - | 448 | $2.14 \pm 0.85$ | $1.29 \pm 0.87$ | $3.18 \pm 0.03$ | $1.98 \pm 1.04$ | $1.44 \pm 0.02$ | $0.83 \pm 0.02$ |
| ResNet-152 | - | 224 | $2.54 \pm 1.43$ | $1.19 \pm 0.91$ | $4.28 \pm 0.16$ | $1.77 \pm 1.22$ | $1.38 \pm 0.05$ | $0.79 \pm 0.02$ |
| ResNet-152 | - | 448 | $2.24 \pm 0.88$ | $1.36 \pm 0.92$ | $3.32 \pm 0.06$ | $2.04 \pm 1.17$ | $1.52 \pm 0.03$ | $0.92 \pm 0.02$ |
| ViT-Small | 16 | 224 | $1.84 \pm 0.73$ | $1.16 \pm 0.57$ | $2.73 \pm 0.05$ | $1.74 \pm 0.50$ | $1.24 \pm 0.02$ | $0.77 \pm 0.02$ |
| ViT-Small | 16 | 448 | $2.76 \pm 1.58$ | $1.24 \pm 0.61$ | $4.69 \pm 0.27$ | $1.84 \pm 0.59$ | $1.48 \pm 0.07$ | $0.84 \pm 0.01$ |
| ViT-Small | 8 | 224 | $2.64 \pm 1.05$ | $1.34 \pm 0.60$ | $3.92 \pm 0.03$ | $1.99 \pm 0.43$ | $1.78 \pm 0.02$ | $0.90 \pm 0.02$ |
| ViT-Small | 8 | 448 | $3.24 \pm 1.27$ | $1.59 \pm 0.86$ | $4.74 \pm 0.37$ | $2.48 \pm 0.72$ | $2.23 \pm 0.24$ | $1.00 \pm 0.02$ |
| ViT-Base | 16 | 224 | $2.80 \pm 1.30$ | $1.34 \pm 0.63$ | $4.37 \pm 0.23$ | $2.01 \pm 0.51$ | $1.75 \pm 0.11$ | $0.90 \pm 0.03$ |
| ViT-Base | 16 | 448 | $2.82 \pm 1.15$ | $1.46 \pm 0.75$ | $4.21 \pm 0.20$ | $2.23 \pm 0.66$ | $1.89 \pm 0.09$ | $0.95 \pm 0.02$ |
| ViT-Base | 8 | 224 | $4.42 \pm 1.78$ | $1.85 \pm 0.81$ | $6.60 \pm 0.08$ | $2.78 \pm 0.46$ | $2.97 \pm 0.02$ | $1.24 \pm 0.02$ |
| ViT-Base | 8 | 448 | $5.26 \pm 2.10$ | $2.15 \pm 0.88$ | $7.82 \pm 0.24$ | $3.10 \pm 0.65$ | $3.56 \pm 0.08$ | $1.52 \pm 0.08$ |
